# Age-dependent Structural Reorganization of the Human Plasma Proteome

**DOI:** 10.64898/2026.07.19.739388

**Authors:** Jaeho Ji, Yeonhee Kim, Eunjeong Han, Yunho Choi, Jumin Park, Heejin Lee, Sujin Park, Ahrum Son, Hyunsoo Kim

**Affiliations:** Graduate School of Life Sciences, Chungnam National University, Daejeon 34134, Republic of Korea; Department of Convergent Bioscience and Informatics, Chungnam National University, Daejeon 34134, Republic of Korea; Graduate School of Medical Science, University of Ulsan, Ulsan 44610, Republic of Korea; Department of Bio-AI Convergence, Chungnam National University, Daejeon 34134, Republic of Korea; Institute of Molecular Life Sciences, Chungnam National University, Daejeon 34134, Republic of Korea; Protein Design Institute, Chungnam National University, Daejeon 34134, Republic of Korea; SCICS, Daejeon 34134, Republic of Korea

**Keywords:** Aging, Plasma Proteome, Structural Proteomics, Covalent Protein Painting, Protein– Protein Interaction

## Abstract

Aging is characterized by progressive physiological decline, yet the molecular mechanisms underlying these changes remain incompletely understood. Although extensive work has examined age-related alterations in protein abundance, the age-dependent structural reorganization of proteins and their interaction networks remains largely unexplored. Here, we quantitatively characterize the structural dynamics of plasma proteins across healthy adult aging and identify age-dependent accessibility patterns that may serve as candidate structural biomarkers. We employed Covalent Protein Painting (CPP) coupled with multiple reaction monitoring (MRM)-based LC-MS/MS to analyze plasma from 24 healthy individuals spanning four age groups (20s to 50s, n = 6 per group). Lysine surface accessibility changes were analyzed using cosine similarity trajectory matching and limma-based linear modeling. The intersection of the two approaches defined five reproducible age-dependent trajectories encompassing 27 proteins and 57 peptides. Integrating STRING/Cytoscape network analysis with AlphaFold3 multimer complex prediction localized several CPP-detected peptides to within ∼5 Å of predicted protein-protein interaction interfaces (KNG1, C3; REN, ALB; and ALB, FGB), suggesting that age-associated accessibility changes are organized at interaction interfaces rather than occurring as isolated local events. Our findings indicate that the human plasma proteome undergoes coordinated, network-level changes in structural accessibility during adult aging, providing a framework for structure-based biomarker discovery. Because CPP signal reflects both conformational accessibility and protein abundance, and because the cohort is modest and cross-sectional, the trajectories reported here are framed as hypothesis-generating and are interpreted with these constraints made explicit throughout.

## Introduction

Aging is the gradual, time-dependent decline in the physiological function of an organism and is the principal risk factor for a broad spectrum of chronic diseases, spanning cancer, cardiovascular and metabolic disease, and neurodegeneration^1^. and is the principal risk factor for a broad spectrum of chronic diseases. A mechanistic understanding of aging is therefore central to preventive and therapeutic strategies for age-related disorders. The influential hallmarks-of-aging framework has organized this complexity into a set of interconnected molecular and cellular processes—among them genomic instability, epigenetic alteration, cellular senescence, and, of central relevance here, the progressive loss of proteostasis. Because these hallmarks are increasingly regarded as tractable points of therapeutic intervention, a mechanistic, molecular-level account of how aging remodels the proteome is central to the development of preventive and therapeutic strategies for age-related disorders.

Molecular studies of aging have historically been anchored in the transcriptome^2^, and numerous analyses have modeled the quantitative relationship between chronological age and gene expression, culminating in transcriptomic “aging clocks” that estimate biological age and distinguish healthy from pathological aging across diverse tissues^3-5^. The maturation of quantitative mass spectrometry subsequently extended this paradigm from RNA to protein, and many contemporary studies interpret the biology of aging primarily through age-dependent changes in protein concentration; in human plasma in particular, large-scale abundance profiling has revealed that the proteome does not drift uniformly but rises and falls in distinct waves across the lifespan, defining an accessible, blood-based readout of systemic aging^6-8^. Powerful as it is, this abundance-centric view captures only one dimension of proteome state—how much of each protein is present—while leaving a second, and equally functional, dimension largely unmeasured.

Protein function, however, is determined not by concentration alone but by three-dimensional structure and by the molecular interactions that structure enables; proteins populate dynamic conformational ensembles whose shifts couple genotype to phenotype, so that two samples with identical protein concentrations may nonetheless differ profoundly in functional state when the underlying conformational or interaction landscape has changed^9,10^. An abundance-only description of the aging proteome is therefore inherently incomplete: it cannot report on folding state, on the exposure or burial of functional surfaces, or on the assembly and disassembly of protein complexes— precisely the structural properties through which proteins execute their biological roles.

These structural properties are not incidental detail; they are actively remodeled during aging. Proteome-wide surveys have shown that discrete, aging-associated conformational transitions accumulate across many proteins^11^. Mechanistically, the proteostasis network that maintains protein folding, stability, and turnover is organized into functional modules that deteriorate progressively with age^12^, and the resulting failures of conformational quality control—misfolding, destabilization, and aggregation—are causally linked to age-related pathology, most conspicuously in neurodegenerative disease^13^. Emerging structure-resolved studies have begun to make this explicit at the level of individual proteins, demonstrating, for example, that the intrinsic capacity of proteins to refold declines with age^14^. Yet work that directly interrogates protein structural change during aging remains scarce relative to the vast literature on abundance and post-translational modification, and almost none extends this structural view to the human plasma proteome or resolves it at the level of protein– protein interaction interfaces—leaving a clear and consequential gap^15,16^.

Human plasma is an especially compelling medium in which to close this gap: it is minimally invasive to obtain, integrates physiological signals from tissues throughout the body, and is already the substrate of choice for clinical biomarker discovery, so a structural readout of the plasma proteome could translate directly toward measures of biological age. Mass spectrometry–based structural proteomics now makes such measurements feasible in complex, native samples, with approaches that read out solvent accessibility or conformational state detecting functional structural alterations at high resolution directly in situ^9,10^. Among these, Covalent Protein Painting (CPP) quantifies the solvent accessibility of lysine ε-amines by reductive dimethylation, providing a peptide-resolution report of protein folding and interaction state: a lysine that becomes buried upon complex formation or compaction loses signal, whereas one that becomes exposed gains it^15,16^. CPP has recently been deployed at proteome scale in living systems and, notably, has resolved plasma-protein structural signatures capable of classifying disease status in Alzheimer’s disease, establishing both its sensitivity in blood and its direct relevance to aging biology^17,18^. Coupling CPP to multiple reaction monitoring (MRM)-based LC-MS/MS adds the targeted specificity and quantitative reproducibility required to track defined accessibility features reproducibly across sample groups.

To convert these measurements into age-structured biology, we integrate targeted quantification with a two-tiered computational framework. Each peptide’s decade-to-decade accessibility trajectory is matched by cosine similarity—a metric well established in tandem mass spectrometry and biological-sequence analysis—against a set of idealized reference patterns, isolating features whose change is directionally age-structured^19,20^. Statistical significance is then established independently with the limma empirical-Bayes linear-modeling framework, which was developed for microarray and RNA-seq data and is now widely applied to quantitative proteomics, and high-confidence features are defined conservatively as the intersection of the two approaches^21-23^. Finally, we exploit the recent transformation of structural biology by deep learning: AlphaFold3-Multimer, combined with STRING/Cytoscape interaction networks, allows us to ask whether the accessibility changes we detect localize to predicted protein–protein interaction interfaces rather than occurring as isolated, local events^24^.

Building on this integrated experimental–computational workflow, our objective is to determine how plasma-protein surface accessibility and interaction patterns are reorganized across healthy adult life (20s–50s) and to nominate candidate structural biomarkers of aging. We analyze plasma from 24 clinically healthy individuals (six per decade, sex-balanced within each group) through the combined CPP–MRM and network/structure-modeling pipeline described above. Throughout, we interpret the findings under two constraints made explicit: the CPP signal convolves conformational accessibility with protein abundance—which we deconvolve using within-protein, lysine-free abundance anchors—and the cohort is modest and cross-sectional, so the trajectories reported here are framed as rigorous, hypothesis-generating candidates rather than established biomarkers.

## Materials and Methods

### Study design and participants

Human plasma was provided by the Ajou University Hospital Human Bio-Resource Bank, and the Kyungpook National University Hospital Human Bioresource Bank, both designated by the Ministry of Health and Welfare. The protocol was approved by the Institutional Review Board (IRB approval no.: 202308-BR-124-01). Participants were assigned to four age groups — 20s, 30s, 40s, 50s — with six individuals per group (total n = 24; mean ± SD age 22.7 ± 1.0, 32.7 ± 1.0, 42.7 ± 1.0, and 52.8 ± 1.3 years, respectively). The cohort was 66.7% female (16 F : 8 M), with an identical sex composition of four females and two males in every decade. Donors were recruited at two biobanks (Ajou University Hospital, n = 9; Kyungpook National University Hospital, n = 15); the two sites were unevenly distributed across decades (the 40s group was drawn entirely from Kyungpook), which we treat as a potential batch/site covariate in the analysis (see Statistical analysis and Discussion). All participants were clinically healthy, with no current smokers, and their baseline clinical laboratory characteristics are summarized in **Table 1**. Consistent with normal physiology, total cholesterol and LDL rose significantly with age (Pearson r = +0.59, p = 0.010 and r = +0.64, p = 0.004, respectively), whereas BMI, fasting glucose, blood pressure, liver enzymes, and complete-blood-count indices showed no significant age trend across this 20s–50s range.

**Table 1.** Clinical characteristics of the study cohort (N=24; mean ± SD or n).

| Characteristic | Overall (N=24) | 20s (n=6) | 30s (n=6) | 40s (n=6) | 50s (n=6) | p (age trend) |
| --- | --- | --- | --- | --- | --- | --- |
| N, n | 24 | 6 | 6 | 6 | 6 | — |
| Age, yr | 37.7 ± 11.5 | 22.7 ± 1.0 | 32.7 ± 1.0 | 42.7 ± 1.0 | 52.8 ± 1.3 | — |
| Female, n (%) | 16 (66.7) | 4 (67) | 4 (67) | 4 (67) | 4 (67) | 1 |
| Site Ajou/Kyungpook | 9/15 | 4/2 | 3/3 | 0/6 | 2/4 | — |
| BMI, kg/m<sup>2</sup> | 22.4 ± 2.9 | 21.9 ± 3.0 | 22.9 ± 4.0 | 23.0 ± 2.9 | 21.7 ± 2.0 | 0.943 |
| SBP, mmHg | 97.4 ± 23.4 | 84.3 ± 29.5 | 93.0 ± 24.6 | 114.8 ± 7.6 | 97.5 ± 20.0 | 0.123 |
| DBP, mmHg | 90.3 ± 20.8 | 98.5 ± 18.2 | 95.7 ± 23.3 | 74.7 ± 8.2 | 92.5 ± 25.2 | 0.276 |
| Glucose, mg/dL | 90.1 ± 13.2 | 84.8 ± 4.2 | 95.2 ± 25.6 | 92.2 ± 6.2 | 88.3 ± 3.2 | 0.643 |
| Total cholesterol, mg/dL | 183.5 ± 31.7 | 161.3 ± 25.5 | 187.2 ± 8.0 | 181.0 ± 29.4 | 215.5 ± 36.0 | 0.01 |
| LDL, mg/dL | 104.2 ± 27.7 | 83.0 ± 14.9 | 104.0 ± 7.1 | 111.8 ± 28.7 | 128.8 ± 36.3 | 0.004 |
| HDL, mg/dL | 61.7 ± 11.8 | 62.8 ± 10.5 | 62.5 ± 14.5 | 51.5 ± 10.5 | 69.2 ± 8.5 | 0.981 |
| Triglyceride, mg/dL | 103.2 ± 83.1 | 77.8 ± 27.9 | 103.2 ± 49.0 | 157.5 ± 171.5 | 86.8 ± 22.0 | 0.478 |
| AST, IU/L | 21.9 ± 7.4 | 18.7 ± 4.2 | 22.2 ± 10.3 | 24.5 ± 7.7 | 22.2 ± 6.8 | 0.261 |
| ALT, IU/L | 18.9 ± 12.2 | 12.7 ± 5.2 | 20.0 ± 12.0 | 25.5 ± 17.6 | 17.5 ± 10.1 | 0.282 |
| GGT, IU/L | 19.3 ± 11.3 | 15.2 ± 5.6 | 18.7 ± 8.8 | 24.0 ± 19.2 | 19.3 ± 7.6 | 0.274 |
| Creatinine, mg/dL | 0.7 ± 0.1 | 0.7 ± 0.2 | 0.7 ± 0.1 | 0.6 ± 0.1 | 0.7 ± 0.1 | 0.477 |
| WBC, $\times 10^3/\mu\text{L}$ | 5.9 ± 1.8 | 6.3 ± 1.4 | 6.6 ± 3.0 | 5.6 ± 0.7 | 5.0 ± 1.0 | 0.172 |
| Hb, g/dL | 13.7 ± 1.8 | 13.0 ± 1.9 | 14.3 ± 1.2 | 13.4 ± 2.5 | 13.9 ± 1.6 | 0.411 |
| Platelets, $\times 10^3/\mu\text{L}$ | 247.4 ± 59.1 | 243.3 ± 29.0 | 253.2 ± 23.4 | 247.5 ± 92.3 | 245.5 ± 77.9 | 0.991 |
| Current smoker, n | 0 | 0 | 0 | 0 | 0 | — |
| Current drinker, n | 5 | 2 | 2 | 0 | 1 | — |

p = Pearson correlation with age (continuous)

### Covalent protein painting (CPP)

CPP was performed to quantify the surface accessibility of plasma-protein lysine residues in each age group. Protein concentration was measured by BCA assay (Pierce, Thermo Scientific). Following the CPP protocol^15,17,18^ solvent-accessible lysine ε-amines were labeled by reductive dimethylation using a solution containing 1% ¹³CD□ O and 300 mM NaBD□CN (medium/heavy dimethyl channel, with shaking at 20 °C for 30 min. The reaction was quenched with 50 mM ammonium bicarbonate (20 °C, 10 min). Proteins were purified by methanol/chloroform precipitation, resuspended in 2% sodium deoxycholate (SDC) with 10 mM TCEP, and denatured at 60 °C for 1 h. Cysteines were alkylated with 20 mM iodoacetamide (dark, 25 °C, 30 min). Because CPP dimethylates lysines — blocking trypsin cleavage at K — digestion used chymotrypsin (37 °C, 16 h), which cleaves independently of lysine labeling and is therefore appropriate for CPP workflows. Digestion was quenched with 1% formic acid; samples were centrifuged and 95 µL of supernatant transferred to LC-MS vials. Immediately before analysis, 5 µL of the 6×5 LC-MS/MS Peptide Reference Mix (Promega) was added to each vial as internal standard (IS).

### Mass spectrometry

CPP-labeled plasma was analyzed on an Agilent 1290 Infinity II UHPLC coupled to an Agilent 6495D triple-quadrupole mass spectrometer with electrospray ionization. Mobile phases were (A) 0.1% formic acid in water and (B) 0.1% formic acid in acetonitrile. Detectability was first assessed on pooled plasma in unscheduled MRM mode to confirm transitions. Method building and data processing were performed in Skyline (v22.2). Decoy transitions (reversed sequences at 50% of the target-transition rate) were included to support mProphet peak scoring. Each unscheduled method contained up to 186 transitions (cycle time 1000 ms). Peaks were integrated with an mProphet scoring model retaining the top three transitions per precursor. Quantification of individual samples (n = 24) then used scheduled MRM (4-min retention window, up to 165 concurrent transitions, three injection methods; injection volume 10 µL, cycle time 1000 ms), processed in Skyline with the same mProphet model at q < 0.05. Integrated peak areas were exported for statistical analysis.

### Quantification and normalization

Peptide-ion peak areas from Skyline were normalized to the IS to yield peak-area ratios (PAR). From the IS reference peptide, the two most intense transitions of each of the top three precursors were used, giving six candidate IS ions (PAR1–PAR6). PAR was computed as (peptide-ion peak area)/(corresponding IS peak area). Reproducibility of each candidate was assessed by coefficient of variation (CV = SD/mean × 100). The log□□(PAR) distribution was inspected as a histogram. Calculations and visualization used R (v4.3.2).

### Statistical analysis

Raw PAR values were log-transformed and z-score standardized (Python), then imported into R. Per-group means (20s, 30s, 40s, 50s) were used for cosine-similarity analysis, which compared each feature’s decade-change vector (Δ30s−20s, Δ40s−30s, Δ50s−40s) to idealized reference patterns whose components take values in <−1, 0, +1> (decrease, stable, increase). Cosine similarity was computed by the standard dot-product formula; the degenerate (0,0,0) template (a zero vector, for which cosine is undefined) was excluded, leaving 26 informative templates. Features with cosine ≥ 0.90 to a given template were considered concordant with that template; because a feature may exceed 0.90 for more than one template, per-template feature sets are not mutually exclusive and are not summed (each feature’s primary assignment is the maximum-cosine template).

For statistical validation, log□(PAR) values were fit with limma (‘lmFit’) using empirical-Bayes moderation (‘eBayes’, ‘trend = TRU’) to account for intensity-dependent variance. Two complementary designs were used: (i) a one-way model across the four decades (moderated F-test for any age effect) and (ii) a continuous age-trend model (slope per year), from which the log□ fold-change across the 20s→50s span and its 95% confidence interval were derived from the moderated standard error. P-values were adjusted for multiple testing by the Benjamini–Hochberg procedure, and significance was defined as adjusted q < 0.05 with |log□FC| > log□(1.5). Because the cohort is exactly sex-balanced within each decade (4 F : 2 M per group), sex is orthogonal to age by design and its inclusion as a covariate does not alter the age contrasts; recruitment site, by contrast, is partially confounded with age (Spearman ρ = +0.41, p = 0.044) and is the more consequential nuisance variable. Each feature’s trajectory was encoded from the sign of consecutive-decade group-mean changes (increase = +1, decrease = −1, stable = 0), and high-confidence age-related features were defined as the intersection of cosine-concordant (≥ 0.90) and BH-significant features. All analyses used R (v4.5.3) with the limma package.

### Network analysis and structural modeling

Protein IDs for pattern-selected features were converted to gene IDs and mapped to STRING within Cytoscape (species: *Homo sapiens*; network: full STRING network; confidence score ≥ 0.7) to build pattern-specific interaction networks.

Protein–protein pairs within the pattern-specific networks were modeled with AlphaFold3-Multimer using UniProt sequences. N-linked glycosylation sites were specified as NAG to represent the PTM. For each complex we report mean pLDDT, ipTM, pTM, and interface-PAE. PyMOL was used to map CPP-identified peptides onto the predicted complexes and to measure peptide-to-interface distances.

## Result

### Selection of an optimal internal standard for normalization

To choose a normalization ion, six candidate IS ions (PAR1–PAR6) were derived from YVYVADVAAK of the Promega 6×5 reference mix, added to every sample before analysis. YVYVADVAAK was selected for its stable intensity and consistent detection across runs; the two highest-intensity transitions (y8, y6) of its top three precursors gave the six candidates. We compared PAR stability and reproducibility across candidates. The log□□(PAR) distribution for each IS was examined for the fraction of values within the target range (−1 ≤ log□□PAR ≤ 1; i.e. 0.1–10 on a linear scale) (**Figure 1A**). PAR3, PAR4, and PAR5 showed the largest in-range fractions (**Figure 1B**): stand1 was lowest (15.4%), while stand4 (53.0%) and stand5 (51.5%) exceeded 50%. Reproducibility, assessed by CV (**Figure 1C**), was best for PAR2 (20.2%) and worst for PAR6 (57.9%); stand5, despite a high in-range fraction, had an elevated CV (39.1%). Stand4 (PAR4 of y6 ion; peptide YVYVADVAAK) combined the highest in-range fraction (53.0%) with an acceptable CV (25.5%) and was selected as the normalization IS for all downstream analyses.

**Figure 1.**
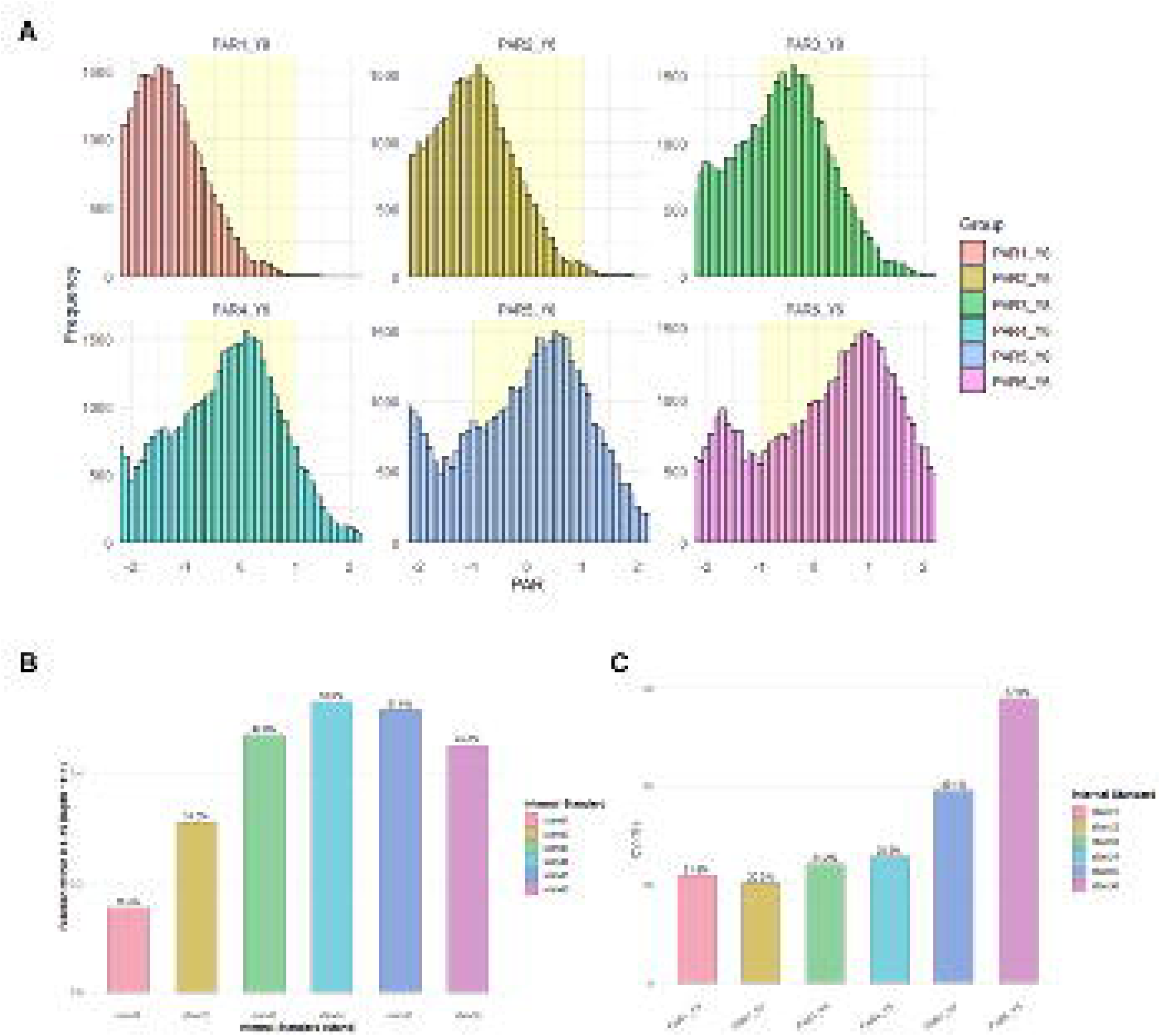
Comparative evaluation of internal-standard normalization performance. (A) Distribution of log□□(PAR) values across the six internal standards (PAR1_y8–PAR6_y6). The yellow-shaded band marks the target range (−1 ≤ log□□PAR ≤ 1), equivalent to 0.1–10 on a linear scale. (B) Proportion of PAR values falling within the target range for each internal standard, with percentages annotated above each bar. Standards 4 and 5 performed best, each retaining >50% of values within the target range. (C) Coefficient of variation (CV%) across internal standards, where lower values reflect greater normalization reproducibility. Standards 5 (PAR5_y8) and 6 (PAR6_y6) showed elevated CVs, indicating greater normalization variability.

### Cosine-similarity trajectory matching identifies age-patterned features

To find features with age-structured accessibility trajectories, we matched each feature’s decade-change vector (Δ30s−20s, Δ40s−30s, Δ50s−40s) against the informative reference templates (components ∈ <−1, 0, +1>; the degenerate (0,0,0) template excluded). Features with cosine ≥ 0.90 to a template were considered concordant. Two representative templates are shown (**Figure 2**). For the (−1, 0, 0) template — an early decrease (20s→30s) followed by stabilization — concordant features (cosine ≥ 0.90) traced exactly this decline-then-plateau trajectory (**Figure 2A**). For the (0, 1, 0) template — mid-life rise (30s→40s) with flanking stability — concordant features rose in middle age and plateaued thereafter (**Figure 2B**). Because per-template concordance sets overlap (a feature can match several templates at ≥ 0.90), they are reported as template-level concordance counts rather than disjoint partitions, and final feature membership is fixed at the intersection step below.

**Figure 2.**
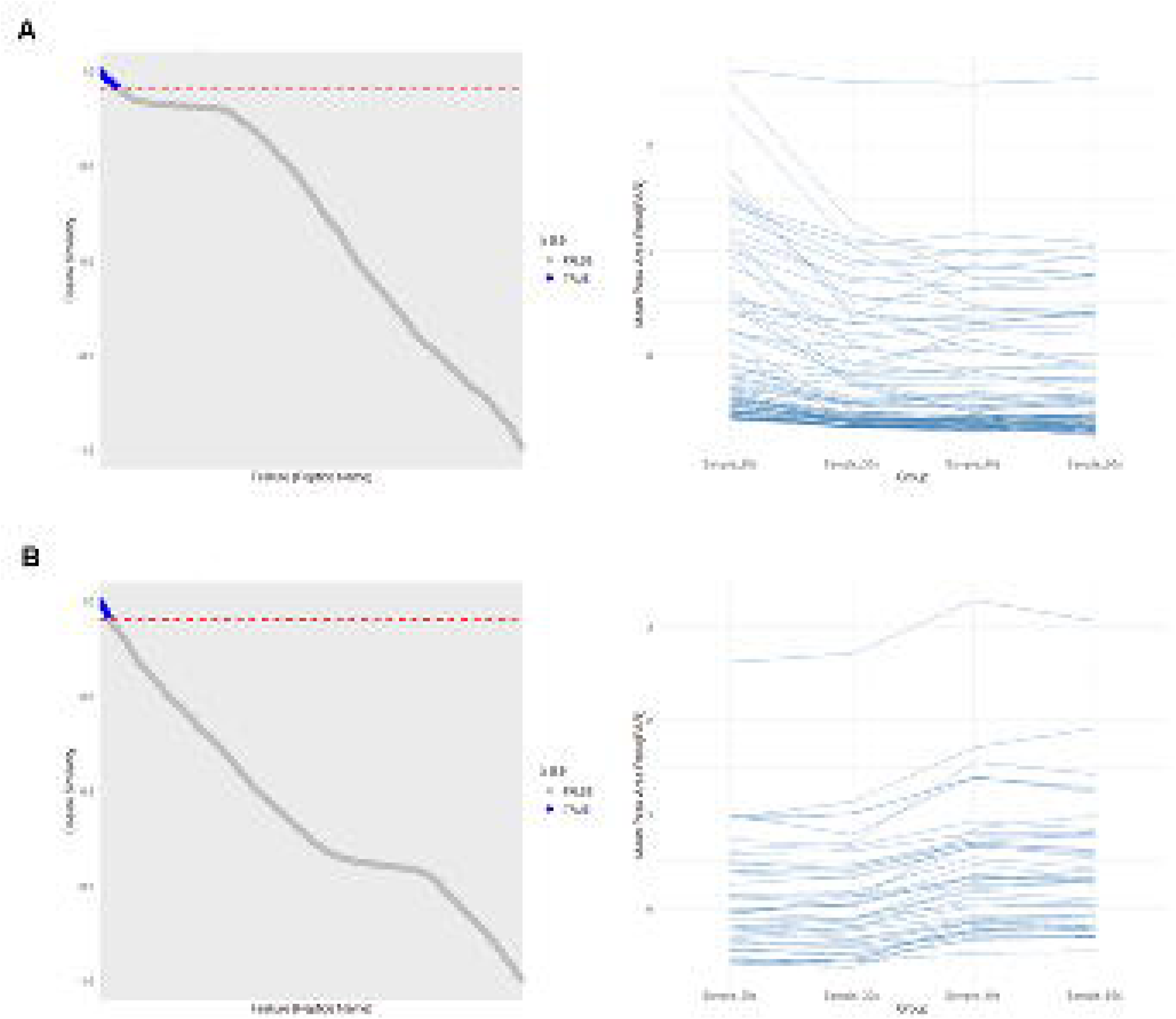
Identification of age-specific ion features by cosine-similarity analysis. (A) Analysis against the reference pattern (−1, 0, 0). Left: distribution of cosine-similarity values across all quantified features; blue points (TRUE) denote features closely matching the reference pattern (cosine ≥ 0.9) and gray points (FALSE) those below the threshold, with the red dashed line marking the 0.9 cutoff. Right: age-dependent trajectories (20s–50s) of representative matched features, showing an initial decline followed by stabilization. (B) Analysis against the reference pattern (0, 1, 0). Left: cosine-similarity distribution. Right: trajectories of pattern-matched features, showing a pronounced mid-life elevation (30s–40s) followed by a plateau.

### Integrating limma modeling with cosine similarity defines five high-confidence trajectories

We applied limma linear modeling to log□(PAR) with empirical-Bayes moderation, using two complementary designs: a one-way model across the four decades (moderated F-test for any age effect) and a continuous age-trend model (slope per year). P-values were adjusted by the Benjamini– Hochberg procedure. The continuous-trend model identified 316 ions from 55 proteins at BH-FDR < 0.05 (**Figure 3A**), reported with log□fold-change over the 20s→50s span and 95% confidence intervals (Table S1); the stricter across-decade F-test retained 25 ions. The importance of correction is direct: an uncorrected p < 0.05 threshold would have declared 403 ions significant by the F-test alone.

**Figure 3.**
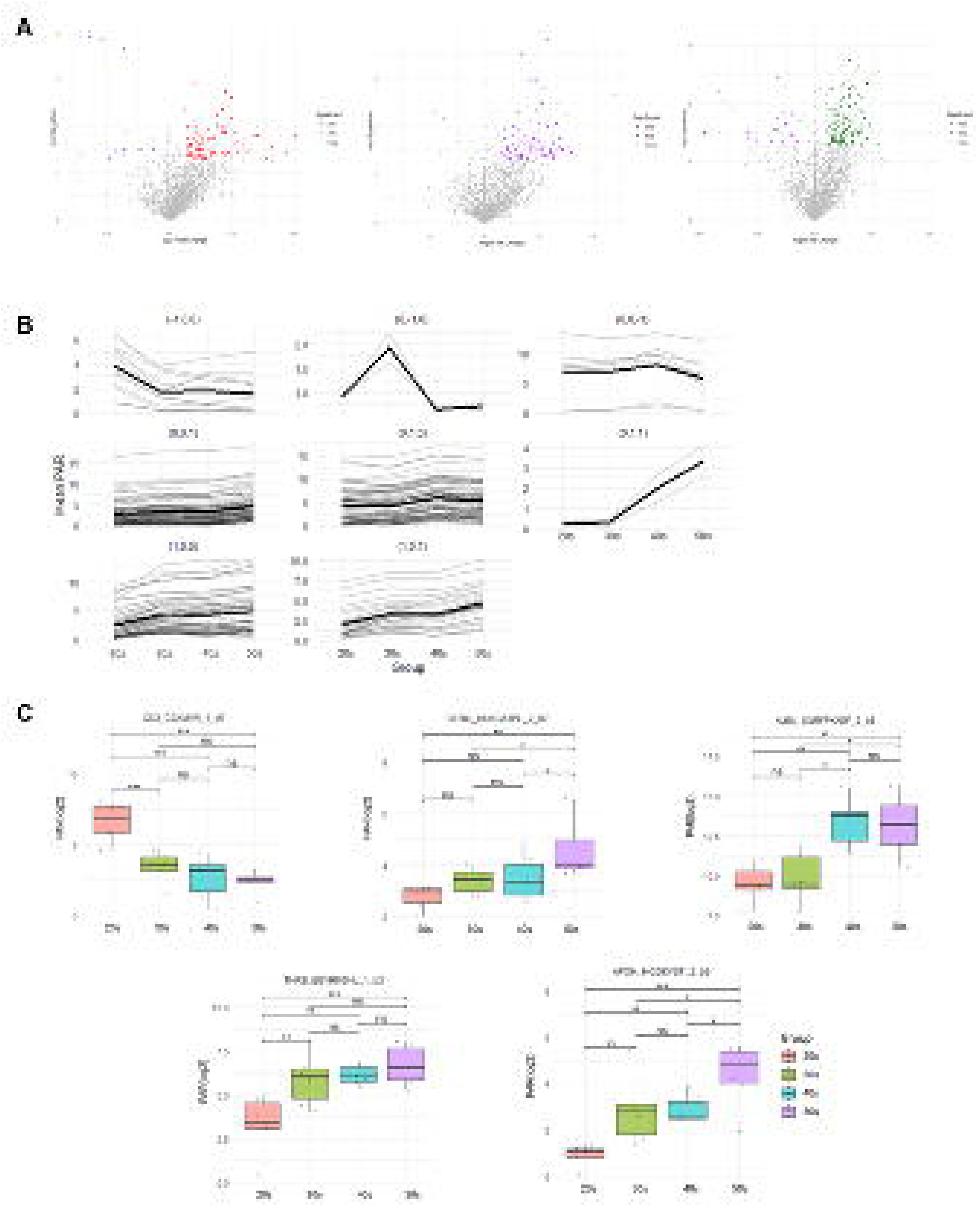
Definition of high-confidence, age-related structural trajectories by integrated limma– cosine analysis. (A) Volcano plots of differential ion abundance between consecutive age groups (20s vs. 30s, 30s vs. 40s, 40s vs. 50s), modeled with limma. Features meeting the significance criteria (p < 0.05 and |log□FC| > log□1.5) are shown in color; non-significant features are gray. (B) Eight distinct age-dependent conformational trajectories derived from the limma analysis. Thin lines trace individual ions; bold lines denote the mean trend of each pattern (1, increasing; 0, stable; −1, decreasing). (C) Five canonical age-associated structural patterns defined by convergent limma–cosine analysis. Boxplots show log□(PAR) values for representative ions from each trajectory. Significance: ns (p ≥ 0.05), * (p < 0.05), ** (p < 0.01), *** (p < 0.001).

A striking feature of the significant set is its directionality: 308 of 316 ions increased with age and only 8 decreased (**Figure 3A**). A near-unidirectional shift affecting ∼16% of all measured ions is difficult to reconcile with 308 independent conformational events and instead points to a global component — internal-standard normalization drift, sample loading, or a systematic change in labeling efficiency — superimposed on protein-specific signal. We therefore treat between-ion normalization as a prerequisite for biological interpretation and address the abundance contribution explicitly below.

By encoding the sign of consecutive-decade group-mean differences (+1 increase, −1 decrease, 0 stable) and matching to reference templates (cosine ≥ 0.90), each ion was assigned to one of the non-constant trajectory classes (**Figure 3B**). Intersecting cosine-concordant assignment with limma significance produced a high-confidence set of 314 ions across 55 proteins. The set is dominated by increasing patterns — (1,0,1) n = 101, (1,1,1) n = 84, (1,1,0) n = 48, (1,0,0) n = 34 — with decreasing classes rare (≤ 3 ions each), consistent with the global upward shift noted above (**Table 2**). For each representative ion, age-specific log□(PAR) values were visualized as boxplots (**Figure 3C**), with pairwise statistical significance denoted as ns, *, **, and *** for p ≥ 0.05, p < 0.05, p < 0.01, and p < 0.001, respectively.

**Table 2.** High-confidence age-dependent trajectory classes (BH-FDR<0.05 & cosine ≥0.90).

| Trajectory (Δ20-30, Δ30-40, Δ40-50) | n ions | n proteins | Pattern |
| --- | --- | --- | --- |
| (1,0,1) | 101 | 27 | ↑20s→30s, ↑40s→50s |
| (1,1,1) | 84 | 26 | ↑20s→30s, ↑30s→40s, ↑40s→50s |
| (1,1,0) | 48 | 25 | ↑20s→30s, ↑30s→40s |
| (1,0,0) | 34 | 14 | ↑20s→30s |
| (0,1,0) | 20 | 12 | ↑30s→40s |
| (0,1,1) | 14 | 6 | ↑30s→40s, ↑40s→50s |
| **(-1,-1,-1)** | 3 | 2 | ↓20s→30s, ↓30s→40s, ↓40s→50s |
| **(0,0,-1)** | 2 | 1 | ↓40s→50s |
| **(0,0,1)** | 2 | 2 | ↑40s→50s |
| **(-1,0,0)** | 2 | 2 | ↓20s→30s |
| **(1,1,-1)** | 1 | 1 | ↑20s→30s, ↑30s→40s, ↓40s→50s |
| **(-1,-1,0)** | 1 | 1 | ↓20s→30s, ↓30s→40s |
| **(0,1,-1)** | 1 | 1 | ↑30s→40s, ↓40s→50s |
| **(-1,0,-1)** | 1 | 1 | ↓20s→30s, ↓40s→50s |

### Abundance-independent accessibility changes isolated using within-protein anchors

Because CPP labels only lysine ε-amines, peptides lacking lysine cannot be painted and report protein abundance rather than accessibility. The dataset contains 134 such lysine-free ions spanning 25 of 98 proteins, providing within-protein abundance anchors. For the 1,059 painted ions belonging to these 25 proteins, we subtracted each protein’s own abundance anchor (mean log□PAR of its lysine-free ions) and re-tested the age trend (**Figure 4**). Of 168 painted ions with a significant raw age trend, only 23 (14%) survived abundance correction, while 145 (86%) were abundance-driven and 13 were newly revealed after removing abundance variance. Thus, for proteins where the correction is possible, the large majority of the raw PAR age signal reflects changing plasma concentration rather than conformational reorganization. The ions that survive anchoring (**Table S2**) constitute the strongest candidates for genuine age-dependent structural change and are the basis for the interface analysis below.

**Figure 4.**
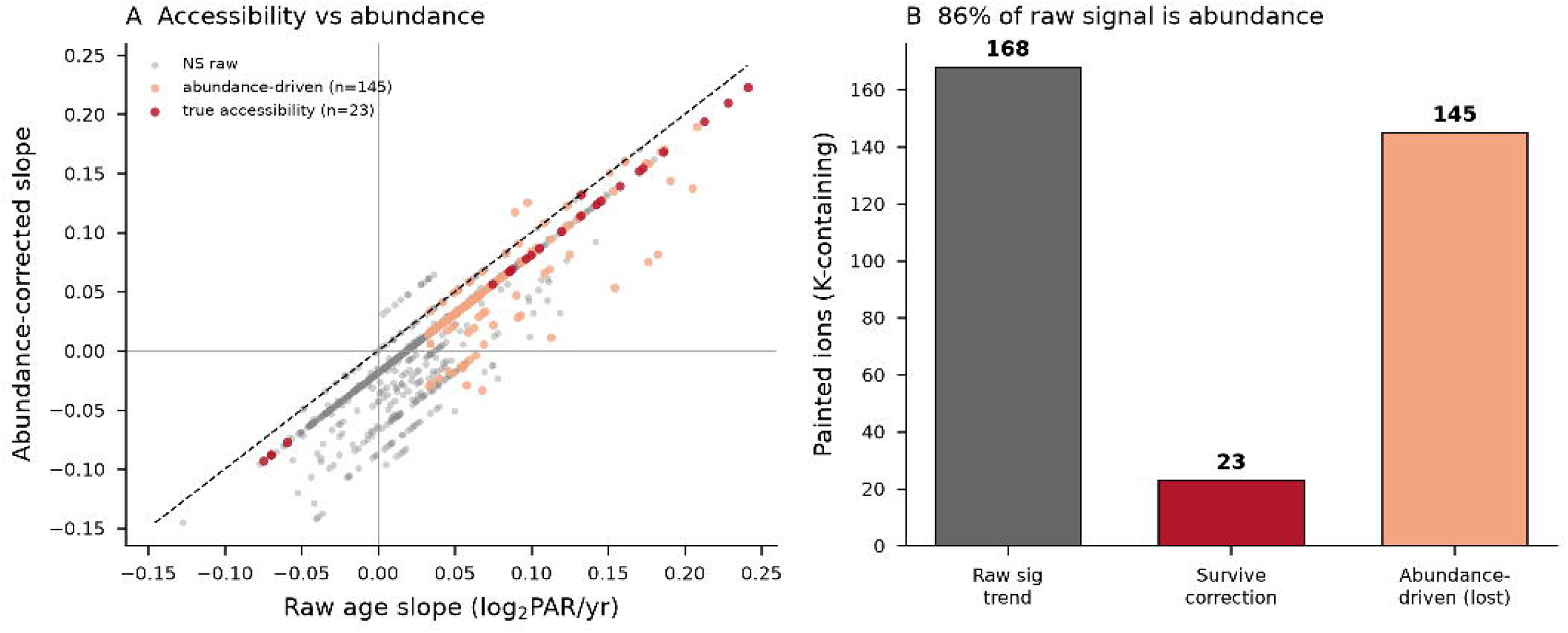
Deconvolving accessibility from abundance using within-protein lysine-free anchors. (**A**) Age-trend slope of each painted ion before (x) versus after (y) subtracting its protein’s abundance anchor (mean log□PAR of lysine-free ions from the same protein); points off the diagonal are driven by abundance. (**B**) Of 168 painted ions with a significant raw age trend, 145 (86%) lose significance after correction (abundance-driven), 23 (14%) survive (abundance-independent accessibility), and 13 are newly revealed. Correction is possible for the 25 proteins carrying an internal lysine-free peptide.

### Clinical characteristics and their relationship to the accessibility signal

Baseline clinical laboratory values for the 24 donors are summarized in **Table 1**. Within this healthy 20s–50s cohort, only lipid measures tracked age: total cholesterol and LDL both rose significantly (r = +0.59, p = 0.010; r = +0.64, p = 0.004), while BMI, blood pressure, fasting glucose, liver enzymes, renal markers, and complete-blood-count indices were age-stable (**Figure 5**). Two design features bear directly on interpretation of the CPP age signal. First, sex is perfectly balanced across decades (4 F : 2 M in each), so the accessibility trajectories cannot be attributed to shifting sex composition. Second, recruitment site is partially confounded with age — the 40s group was drawn entirely from Kyungpook, the 20s predominantly from Ajou (Spearman ρ = +0.41, p = 0.044) — and the two sites reported certain laboratory indices in different units, indicating distinct pre-analytical pipelines. Because the CPP age signal is near-unidirectional (308/316 ions increasing), site/batch is a plausible contributor alongside abundance and normalization drift, and we flag it explicitly as a limitation.

**Figure 5.**
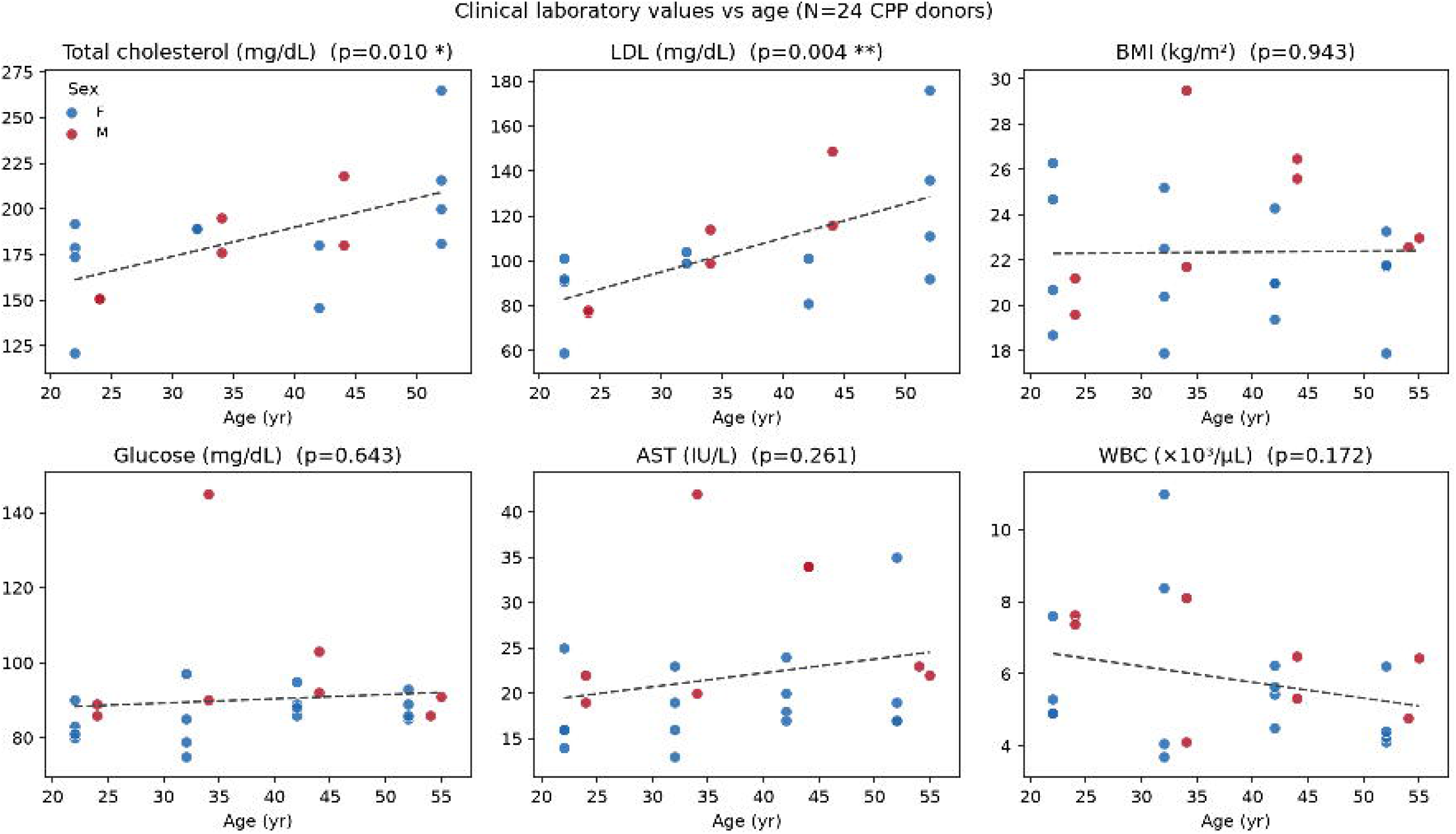
High-confidence age-dependent accessibility trajectories. Group-mean log□(PAR) trajectories (centered on the 20s) for the six most populated trajectory classes among the 314 high-confidence ions (55 proteins; BH-FDR < 0.05 ∩ cosine ≥ 0.90). Class labels give the consecutive-decade sign pattern (Δ30–20, Δ40–30, Δ50–40; +1/0/−1). Increasing patterns dominate — (1,0,1) n = 101, (1,1,1) n = 84, (1,1,0) n = 48, (1,0,0) n = 34 — consistent with the global upward shift in Figure 2.

### Network and structural modeling localize accessibility changes to interaction interfaces

To connect these trajectories to molecular interactions, we built pattern-specific STRING/Cytoscape networks (confidence ≥ 0.7, Homo sapiens) and modeled selected protein pairs with AlphaFold3-Multimer, specifying N-glycosylation sites (NAG). Recognizing that CPP reports labeling of solvent-exposed lysine ε-amines — so that burial of a lysine upon complex formation reduces its PAR, and exposure increases it — we asked whether CPP-detected peptides sit near predicted interfaces.

Three points require explicit statement so the structural claims are not over-read. First, the APOH peptide HGDKVSF showed the most robust age dependence in the entire dataset (log□FC ≈ +3.3 to +3.7; BH-FDR 0.019–0.038 by F-test, 0.004–0.008 by trend), and the albumin SQRFPKAEF and complement C3 (EKQKPDGVF, RNNNEKDMAL) peptides were trend-significant (BH-FDR < 0.04); these are the defensible structural signals. Second, renin (REN) was not measured in this dataset — no REN peptides are present — so the REN–albumin association is a network/AF3 prediction anchored only on the albumin peptide, and REN is described here as an inferred, not observed, partner. Third, the KNG1 peptide TESCETKKL, the basis of the previously proposed KNG1–C3 (−1,0,0) trajectory, did not survive multiple-testing correction (best-ion BH-FDR = 0.08) and is therefore reported as suggestive only (**Figure 6A**).

**Figure 6.**
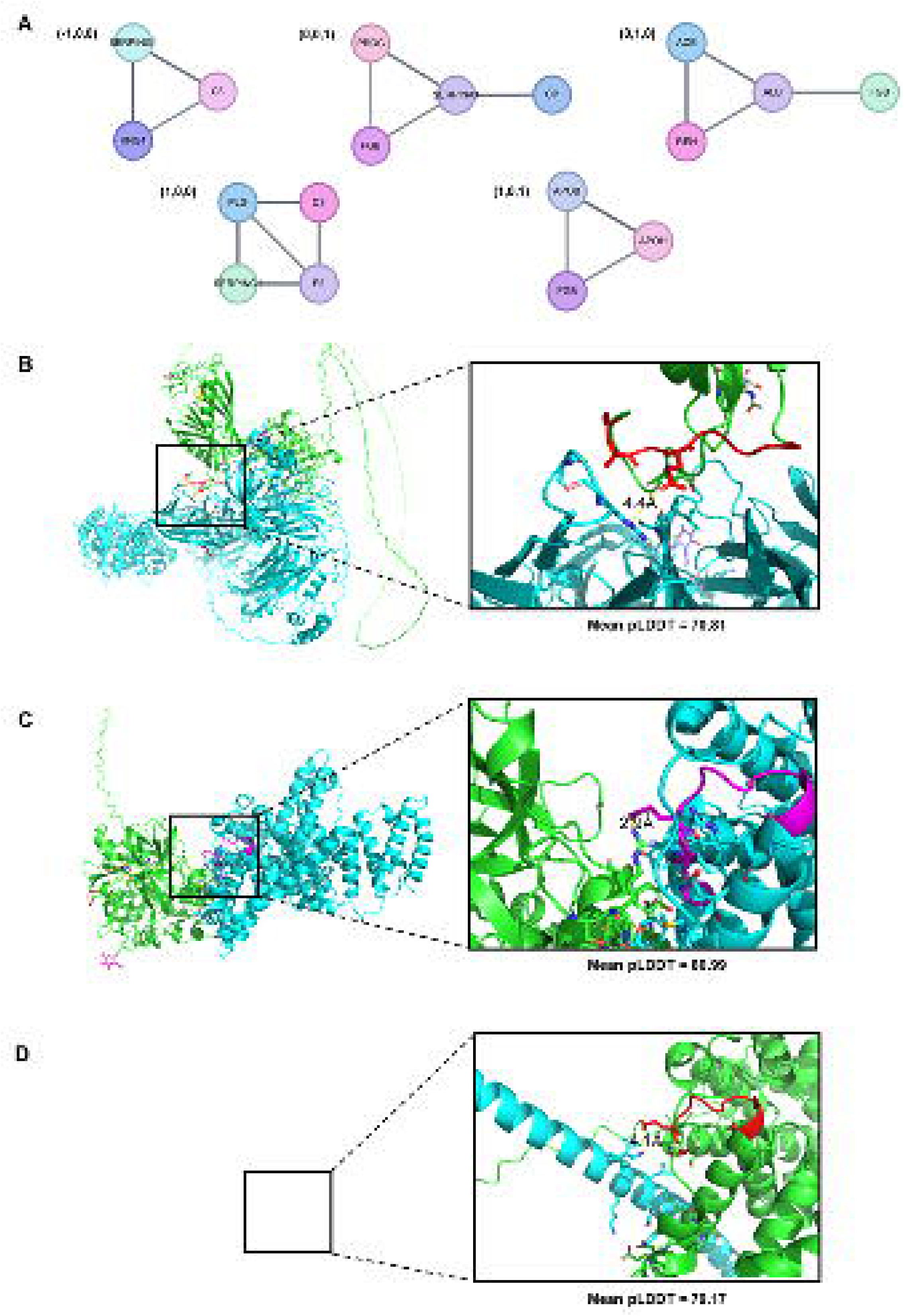
Age-dependent remodeling of protein interfaces revealed by pattern-specific network analysis and AlphaFold3 structural modeling. (A) Pattern-specific protein–protein interaction networks across the aging trajectories, visualized in Cytoscape using the STRING database (confidence ≥ 0.7, Homo sapiens). Each subnetwork corresponds to one of the five canonical patterns: (−1, 0, 0), (0, 0, 1), (0, 1, 0), (1, 0, 0), and (1, 0, 1). (B–D) AlphaFold3-predicted protein complexes exemplifying selected patterns: (B) KNG1–C3 (pattern −1, 0, 0); (C) REN–ALB (pattern 0, 1, 0); and (D) ALB–FGB (pattern 0, 1, 0). Protein chains are color-coded (green, primary protein; cyan, secondary protein), and CPP-identified peptides are highlighted in red/magenta at the interaction interfaces (2.9–4.4 Å). Mean pLDDT scores were 70.8 (B), 81.0 (C), and 79.2 (D).

For the KNG1–C3 complex (pattern (−1,0,0)), the CPP peptide of KNG1 (TESCETKKL, residues 337–345) lay ∼4.4 Å from the C3-binding surface (**Figure 6B**), consistent with a lysine that becomes progressively buried — an early accessibility decrease followed by a maintained, closed interface from the 30s onward. For REN–ALB (pattern (0,1,0)), the ALB peptide SQRFPKAEF (residues 244– 252) lay ∼2.9 Å from the REN-binding surface (**Figure 6C**); the same ALB peptide lay ∼4.1 Å from the interface in the ALB–FGB complex (**Figure 6D**), consistent with mid-life interface opening. Mean pLDDT was 70.8 (B), 81.0 (C), and 79.2 (D). **(Table 3)**

**Table 3.**
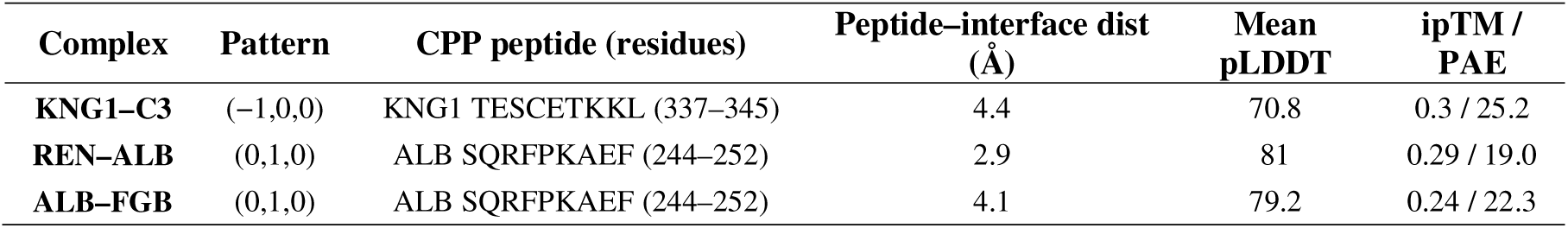
AlphaFold3-Multimer complexes and CPP peptide localization.

Together these results are consistent with age-dependent accessibility changes being organized at protein–protein interaction interfaces rather than as isolated local events, with CPP-detected peptides mapping near predicted binding surfaces. We emphasize these interface assignments are model-based predictions requiring experimental confirmation.

## Discussion

We combined CPP with MRM-based LC-MS/MS to quantify age-dependent changes in plasma-protein lysine accessibility across the 20s–50s, defined reproducible trajectories by intersecting cosine-similarity matching with limma modeling, and used STRING networks with AlphaFold3-Multimer models to place several changes near predicted interaction interfaces. The data are consistent with coordinated, network-level accessibility remodeling during adult aging rather than isolated per-protein changes.

Cosine similarity captures the direction of decade-to-decade change but performs no inference; limma supplies significance, and the intersection yields five high-confidence trajectories. Critically, CPP PAR reflects the product of protein abundance and per-lysine labeling efficiency. A PAR trajectory can therefore arise from a genuine conformational/interface change or from an age-dependent shift in the protein’s plasma concentration — and many of the proteins here (complement C3, kininogen, apolipoproteins, albumin) have known age-associated abundance changes. We addressed this confound directly using the lysine-free peptides already present in the data as within-protein abundance anchors (25 proteins, 1,059 correctable painted ions). The result is sobering and clarifying in equal measure: ∼86% of the raw age signal in anchor-testable proteins is attributable to abundance, and only ∼14% survives as abundance-independent accessibility change. Combined with the near-unidirectional increase of the raw signal (308/316 ions up), this indicates that the unnormalized PAR trajectories substantially overstate conformational reorganization. We therefore restrict genuine structural claims to anchor-surviving ions and present the broader trajectory catalog as accessibility-or-abundance trajectories.

In the AF3 models (KNG1–C3, REN–ALB, ALB–FGB) the CPP peptides (KNG1 TESCETKKL; ALB SQRFPKAEF) fell within ∼5 Å of predicted interfaces, consistent with the measured accessibility directions: the (−1,0,0) KNG1–C3 interface behaves as if it closes early and stays closed, whereas the (0,1,0) REN–ALB/ALB–FGB interfaces behave as if they open in middle age. This peptide-level, interface-centric view is complementary to, and distinct from, the abundance-centric aging-proteome literature [8]. However, AF3 models are static and the interface-confidence metrics (especially ipTM/PAE, not yet reported) must accompany these claims; the KNG1–C3 model in particular is of moderate confidence.

Limitations. (1) The cohort is small (n = 6/decade) and cross-sectional — different individuals per decade — so inter-individual heterogeneity, not longitudinal change, underlies the contrasts. Sex is balanced within each decade (4 F : 2 M), which removes sex as a confounder but leaves the study underpowered to detect sex-specific effects. (2) Recruitment site is partially confounded with age (the 40s group is entirely from one biobank), and the two sites used different pre-analytical conventions; because the CPP age signal is near-unidirectional, an unmeasured site/batch effect is a live alternative explanation that a site-adjusted or batch-corrected reanalysis should address. (3) The abundance/accessibility confound above. (4) The age range stops in the 50s, so conclusions pertain to early-to-mid adult aging, not late-life aging. (5) The normalization anchor’s CV (25.5%) is high; robustness to alternative normalization should be shown. (6) No orthogonal or independent-cohort validation is presented. Future work should couple CPP with an abundance-controlled metric, add an orthogonal structural readout (e.g. LiP-MS) or known structures, incorporate cryo-EM or dynamics where available, and validate candidate structural biomarkers in a larger, longitudinal, sex-balanced cohort.

By integrating accessibility-based structural proteomics (CPP-MRM) with cosine-similarity trajectory matching, limma modeling, STRING networks, and AlphaFold3-Multimer prediction, we show that the human plasma proteome exhibits coordinated, patterned changes in lysine accessibility across healthy adult aging, several of which localize near predicted protein–protein interaction interfaces. These accessibility trajectories are candidate structural signatures of aging and establish a workflow linking targeted quantitative proteomics to structural interpretation. Because CPP signal convolves conformation with abundance and the cohort is modest and cross-sectional, the trajectories are hypothesis-generating; abundance-controlled measurement and validation in larger longitudinal cohorts will be required to establish their biological and biomarker significance.

## Data and code availability

The mass spectrometry data have been deposited to Panorama Public (https://panoramaweb.org/), and are accessible at https://panoramaweb.org/BjjQrY.url. All analysis scripts used in this study, including cosine similarity calculation, differential abundance analysis (limma), and figure generation, are publicly available on GitHub (https://github.com/kimlab-cnu/AgeStructMarker)

## Acknowledgement

The biospecimens and data used for this study were provided by the Biobank of Ajou University Hospital and Korea-Kyungpook National University Hospital (KNUH), a member of the Korea Biobank Network. All materials derived from the Biobank of Korea were obtained (with informed consent) under institutional review board (IRB)-approved protocols. We acknowledge the use of Claude Opus 4.8 solely for linguistic refinement and grammatical corrections in manuscript preparation. All scientific content, data analysis, and intellectual contributions presented herein were developed independently by the authors without the use of generative AI tools.

## Author Contributions

J.J. and Y.K.: conceptualization, methodology, experimental analysis, data analysis, visualization; E.H., Y.C., J.P., H.L., and S.M., : data analysis; J.J., Y.K., A.S, and H.K.: writing-original draft, conceptualization, project administration, resources, supervision, writing-review & editing. All authors have read and approved the final manuscript.

